# MicroED structure of lipid-embedded mammalian mitochondrial voltage dependent anion channel

**DOI:** 10.1101/2020.09.17.302109

**Authors:** Michael W. Martynowycz, Farha Khan, Johan Hattne, Jeff Abramson, Tamir Gonen

## Abstract

A near-atomic resolution structure of the mouse voltage dependent anion channel (mVDAC) is determined by combining cryogenic focused ion-beam (FIB) milling and microcrystal electron diffraction (MicroED). The crystals were grown in a viscous modified bicelle suspension which limited their size and made them unsuitable for conventional X-ray crystallography. Individual thin, plate-like crystals were identified using scanning electron microscopy (SEM) and focused ion-beam (FIB) imaging at high magnification. Three crystals were milled into thin lamellae. MicroED data were collected from each lamellae and merged to increase completeness. Unmodelled densities were observed between protein monomers, suggesting the presence of lipids that likely mediate crystal contacts. This work demonstrates the utility of milling membrane protein microcrystals grown in viscous media using a focused ion-beam for subsequent structure determination by MicroED for samples that are not otherwise tractable by other crystallographic methods. To our knowledge, the structure presented here is the first of a membrane protein crystallized in a lipid matrix and solved by MicroED.

## Introduction

Determining the crystal structures of membrane proteins embedded in lipids is challenging. A bottleneck in this process for traditional X-ray crystallography is growing large, well-ordered crystals that incorporate ordered or semi-ordered lipids. In contrast to soluble proteins, membrane proteins have both hydrophobic and hydrophilic regions on their surface, making them difficult to handle. To overcome this, membrane proteins are frequently solubilized with detergent and then crystallized but during this process the lipids, that often play critical roles in the structure and function of membrane proteins, are lost (Hunte and Richers, 2008). For this reason several methods were developed for crystallizing membrane proteins within a lipid matrix, for example the lipidic cubic phase and bicelle crystallization (Cherezov, 2011; Faham and Bowie, 2002; Landau and Rosenbusch, 1996; Ujwal and Abramson, 2012). Viscous media such as lipids or lipid/detergent mixtures are used to mimic the hydrophobic environment of a lipid bilayer. This viscous media cannot be easily blotted away using traditional cryoEM blotting methods. Several attempts have been made to circumvent the blotting method by only depositing nano liter volumes by pin printing (Ravelli et al., 2020), vacuuming away the excess media using a pressure differential (Zhao et al., 2019), using liquid wicking grids (Tan and Rubinstein, 2020), and changing the phase of the media using other less viscous detergents (Zhu et al., 2020a). These approaches have been successful in many cases for soluble proteins. However, the forces involved could either dehydrate the membrane protein crystals or permanently damage the lattice. It is preferable to leave membrane protein crystals in their mother liquor and freeze them as quickly as possible to best preserve their hydration and crystalline order.

Prior reports of membrane protein structures determined using MicroED were conducted on crystals of the Ca^2+^ ATPase (Yonekura et al., 2015) and the non-selective ion channel, NaK (Liu and Gonen, 2018). Thin 3D crystals of Ca^2+^ ATPase were grown by dialysis of isolated protein against detergent-free buffer to slowly remove excess detergent. As the detergent was removed ordered layers of lipid-Ca^2+^ ATPase formed and slowly began to stack resulting in thin 3D crystals. These crystals were only a few layers thick making them suitable for analysis by MicroED but they represent a special case where 3D crystals formed serendipitously from stacked 2D crystals. Moreover, this crystallization method required large amounts of purified protein and crystal growth was slow and laborious. In sharp contrast, microcrystals of NaK were identified in conditions using detergents out of a sparse matrix. The crystals formed as small cubes containing only ∼1000 diffracting units. Because the crystals grew out of a detergent solution, without any lipids, they were easy to pipette and excess solution blotted easily for cryoEM grid preparation using standard vitrification equipment (Dubochet et al., 1985). In both examples, the resulting crystals were thinner than ∼500nm so sample preparation for MicroED including data collection were straight forward. However, it is preferable to study membrane proteins with lipids rather than in detergent (Jiang and Gonen, 2012). Crystallization of membrane proteins in lipidic cubic phase or in lipid bicelles regularly yield crystals in the 2-5µm range and these are too small for traditional X-ray crystallography and are too large for MicroED. Moreover, the thick lipid matrix renders such samples almost impossible to prepare by traditional blotting methods for cryoEM because the lipid matrix is viscous and integrated into the crystal. If blotting is inefficient the sample becomes too thick for the electrons to penetrate thus eliminating the ability to perform a MicroED experiment. Strategies for diluting the thick lipid matrix around crystals were recently described using soluble proteins but when applied to membrane proteins the quality of the crystals rapidly deteriorated resulting is low resolution data at best (Zhu et al., 2020b).

The voltage dependent anion channel, VDAC, is a mammalian membrane protein that resides on the mitochondrial outer membrane. Traditionally, VDAC crystals have been studied in both detergent (Bayrhuber et al., 2008; Meins et al., 2008) and within a lipid matrix (Choudhary et al., 2014; Schredelseker et al., 2014; Ujwal et al., 2008) by X-ray crystallography with and without bound ATP. The 31 kDa polypeptide begins with a short N-terminal α-helix that is surrounded by 19 β-strands that pack against one another forming a large barrel-like structure surrounding a hydrophilic pore. The pore of the channel allows cargo up to 5kDa in size to traverse the mitochondrial membrane. Further analyses have identified neurosteroid and cholesterol binding sites (Cheng et al., 2019) and unique roles for individual amino acids and the corresponding lipidic environment (Bergdoll et al., 2018; Betaneli et al., 2012; Rostovtseva and Bezrukov, 2008). Crystallization efforts aimed at determining the structure of mutant mVDAC protein in a modified lipid matrix were stymied because only small crystals, almost indistinguishable by eye, were obtained and efforts to increase their size for traditional X-ray crystallography were unsuccessful, making this an ideal specimen for investigation by MicroED.

Samples in MicroED are prepared similarly to other modalities of cryoEM (Nannenga et al., 2014b; Shi et al., 2016). Briefly, a small amount of liquid is added to a TEM grid, the grid is gently blotted to remove the excess solvent, vitrified in liquid ethane, and stored in liquid nitrogen for future investigation. However, only crystals that happened to be very thin (<500nm) or could be fragmented by sonication or vortexing were amenable to MicroED data collection strategies (de la Cruz et al., 2017; Martynowycz et al., 2017). Unfortunately, membrane protein crystals rarely form crystals thin enough for MicroED. These crystals tend to be more delicate and do not survive the harsh sonication/vortexing treatment so new sample preparation methods are required. Recently, the use of a Focused-Ion Beam on a Scanning Electron Microscope (FIB-SEM) was demonstrated as a viable sample preparation technique for MicroED (Duyvesteyn et al., 2018; Martynowycz et al., 2019a, 2019b) (Figure 1). With this method, an SEM is used to identify crystals and a FIB is used to thin the crystals to <500nm thickness creating the ideal samples for MicroED. Recently two reports, one published and one posted online, attempted to determine the MicroED structure of a membrane protein embedded in LCP but were not successful likely because the sample preparation methods were not optimized (Zhu et al., 2020b; Polovinkin et al., 2020). Importantly, to date FIB milling coupled with successful structure determination by MicroED has only been demonstrated for soluble standard proteins, such as lysozyme and proteinase K (Beale et al., 2020; Duyvesteyn et al., 2018; Li et al., 2018; Martynowycz et al., 2019a, 2019b; Wolff et al., 2020; Zhou et al., 2019).

**Figure 1.**
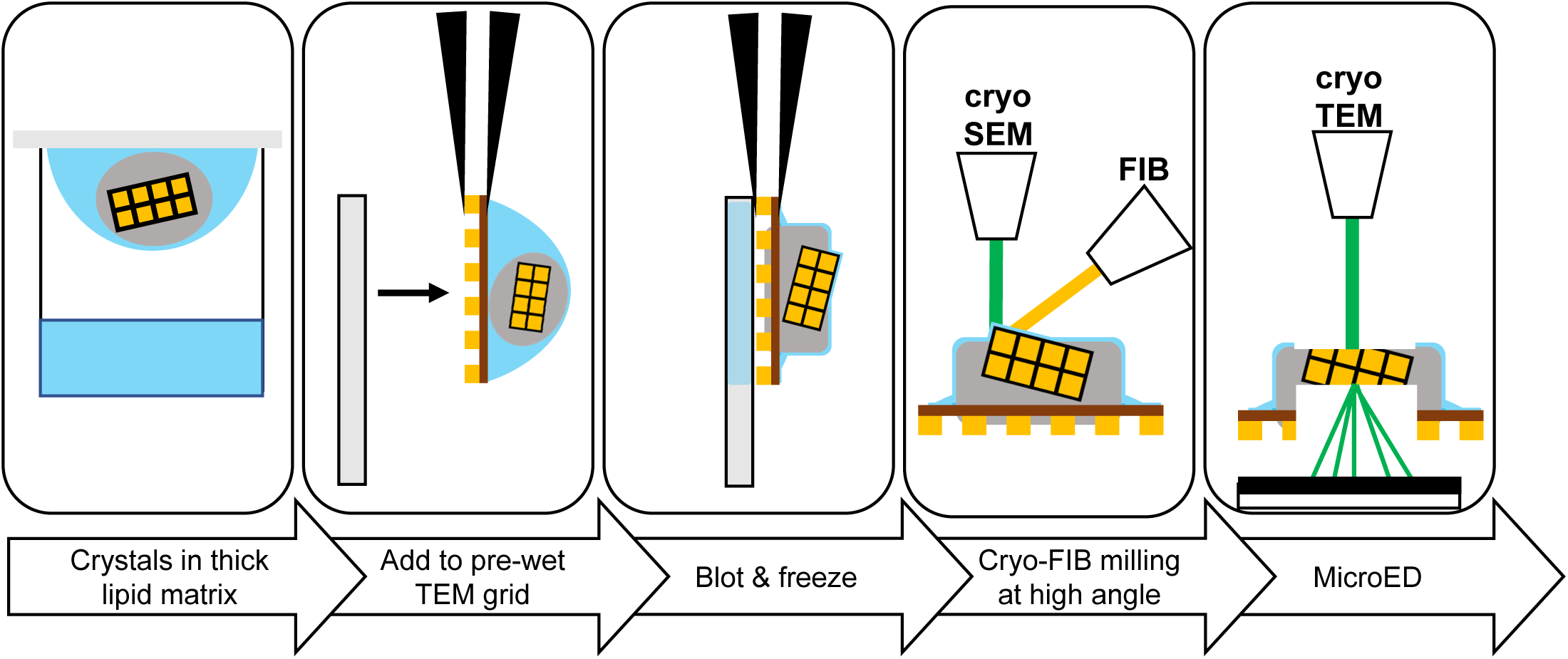
Preparing viscous samples for cryoFIB and MicroED. Schematic cartoon demonstrating the steps in the pipeline for studying membrane protein crystals grown in a lipid matrix by MicroED. Crystals are grown in viscous media, the crystals are transferred in the viscous media to a wet TEM grid, the grid is blotted and the thick media is left behind. Blotted grids are transferred into a FIB/SEM for inspection and milling, and finally thin lamellae prepared by FIB milling are used for MicroED data collection.

Here, we demonstrate the application of FIB milling on membrane protein crystals of mVDAC grown in modified lipid bicelles. Three mVDAC crystals embedded in thick bicelle media were located in the FIB/SEM and milled into ∼200nm thick lamellae. MicroED data were collected and merged to 80% completeness at 3.1 Å resolution. The structure of mVDAC was solved by molecular replacement using a wild type VDAC model. Evidence for lipid packing between mVDAC monomers was visible in the density map. Our results demonstrate that membrane protein crystals embedded in dense, viscous media can be made amenable to MicroED investigation by cryoFIB milling, increasing the scope of the method to include more challenging and biologically important membrane proteins grown in a lipid matrix.

## Results and Discussion

Crystals of mVDAC were grown in modified bicelles in a similar manner as previously described (Ujwal et al., 2008) with subtle changes to lipid composition. In this condition, only very small and thin plate-shaped microcrystal shards were formed within a dense core of a thick and viscous lipid matrix (Figure 2, inset). These microcrystals were difficult to harvest from inside the dense bicelle core in the drop to attempt traditional X-ray crystallography and despite significant effort could not grow larger for analysis by X-ray crystallography. We therefore explored the use of MicroED.

**Figure 2.**
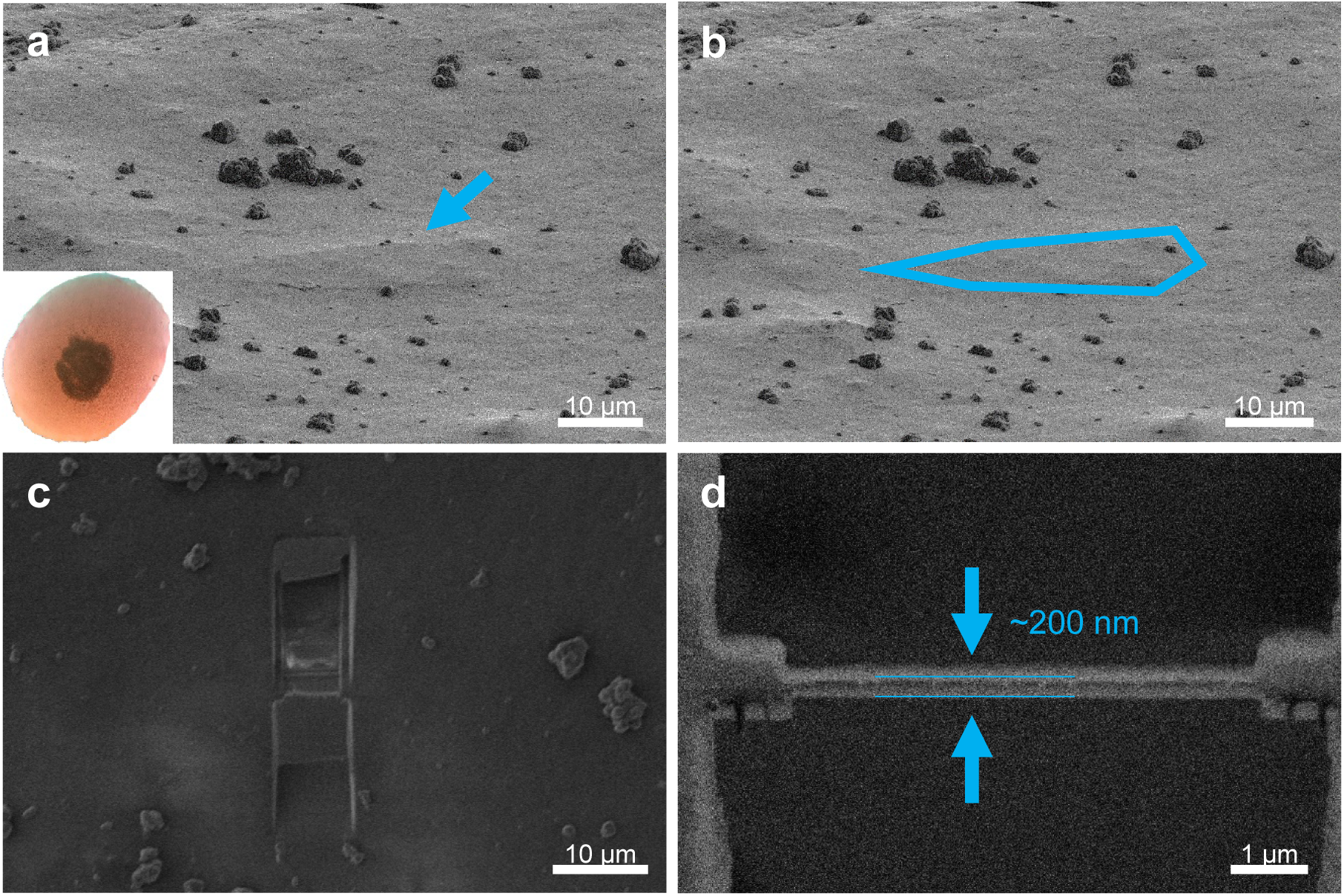
Identification of mVDAC microcrystals in the lipid matrix. (a) FIB image of an identified crystal prior to milling (arrow), and (b) this same picture with the thin crystal outlined in blue. (c) SEM image of the final crystalline lamellae milled at 30° showing stratification between the deposited platinum layer, crystal, and thick bicelle media. (d) FIB image of the final lamellae in the FIB used to measure thickness. mVDAC crystals were ∼200nm thick. Lines indicate top and bottom of lamella. The crystallization drop with bicelle solution is shown inset in (a).

Preparing grids of mVDAC for MicroED analyses proved to be challenging because of the lipid-matrix in the crystallization drops. These were viscous and difficult to handle. The steps we took for optimizing sample preparation of mVDAC crystals in the lipid matrix for MicroED are documented in Supplementary Figures 1-9. With every step the grid preparation improved and the resulting data that was obtained likewise improved leading to high quality data (Table 1) that had high signal to noise ratio (Supplementary Figure 9). We first attempted to prepare grids for MicroED by depositing an entire crystal drop onto the TEM grid and gently blotting from the opposite side as described (Martynowycz and Gonen, 2020). These attempts resulted in samples that were completely opaque and could not be penetrated by the electron beam. At this point we decided to explore the use of a FIB SEM to thin the crystalline material and its surrounding to thicknesses that are ideal for MicroED. Maintaining the grids in a humidity chamber held at ∼35% during blotting resulted in grids with large amorphous material and no visible crystals. Blotting from the front and the back typically broke the windows on the grid and all crystalline material was lost. Grids with less material but visible crystal edges poking through piles of amorphous material were identified by adding mother liquor to the grid prior to blotting at ambient humidity and then rinsing the grids with additional mother liquor and blotting a second time. This double washing step allowed us to identify crystalline material but mostly led to poor or no diffraction so we surmise that the crystalline lattice was damaged by the harsh treatment.

**Table 1.**
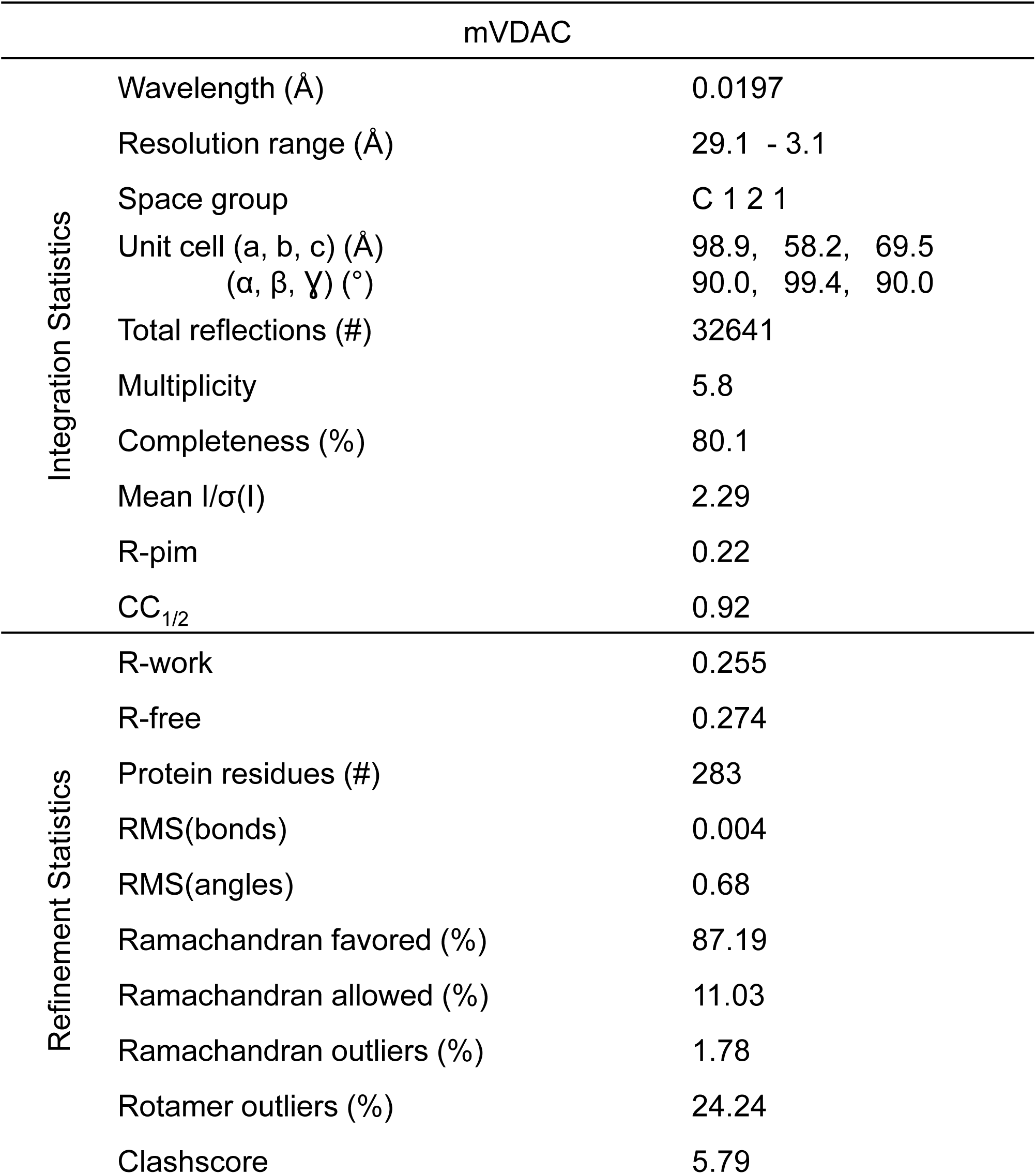
MicroED Crystallographic table.

The breakthrough for MicroED sample preparation occurred when the samples were kept hydrated. We added additional mother liquor on top of the crystal drops to reduce evaporation and minimize lipid exposure to environmental air. In addition, the cryoEM grid was wetted with 2µL of mother liquid inside of a vitrification robot operating at room temperature and 90% humidity and 4°C to prevent any dehydration during sample application. Approximately 0.5µL of the central crystal/lipid matrix was carefully pipetted onto the wet grid. The sample was incubated for 20s and then gently blotted—from behind—and immediately plunged into super cooled liquid ethane for vitrification. Grids were transferred under cryogenic conditions for further investigation using FIB-SEM. Grids with mVDAC crystals were inspected by SEM. Upon Visual inspection, the grid appeared to be covered with a thick layer of material which impeded crystal identification (Figure 1a). Upon careful inspection at high magnification, thin mVDAC crystals were observed under the thick lipid matrix (Figure 1a, arrow and b, outline).

Milling the mVDAC crystals was conducted similarly to prior reports (Martynowycz et al., 2019a, 2019b) with important modifications. We noticed that the viscosity and thickness of the lipid matrix afforded considerable challenges for milling the crystals while minimizing radiation damage or destruction of the underlying crystalline lattice. To determine the total thickness of the preparation, we first milled trenches in front and behind the target crystal. These trenches were milled at a high angle of 35°, compared with the typical milling angle of ∼12°. The high milling angle facilitated the examination of the specimen to decide on milling trajectory and strength (Figure 1c). After trenching, three mVDAC crystals were milled at a maximum angle of 30°. The milling angle was adjusted to assure the crystals would be accessible after rotating the grid for inspection in the TEM. Milling was conducted in sections either above or below the crystal for short periods of time and low currents to prevent overheating the sample and damaging the lattice. We observed that even short bursts of higher currents would over-expose the crystals, destroying the underlying lattice and resulting in little or no diffraction. This observation is in sharp contrast with crystals of soluble proteins that appear to withstand much greater currents and still provide atomic resolution data (Martynowycz et al., 2019b). Therefore, to minimize radiation damage to membrane protein crystals, the ion-beam current was ramped down in steps as the milling progressed (Martynowycz et al., 2019a, 2019b). After a polishing step a thin ∼200nm crystalline lamellae remained (Figure 1d) and the preparation was transferred to a cryo TEM for MicroED.

## MicroED data collection, analysis and structure determination

The milled lamella were loaded onto a 300kV Titan Krios equipped with a CMOS CetaD. Lamella were easy to identify in the TEM. They appeared as a bright stripe against an otherwise black background (Figure 3a). Five hydrated mVDAC lamellae were tested in the Titan Krios, and, to our surprise, 60% of the lamellae diffracted well to approximately 3Å resolution (Figure 3a). Even more surprising, we were able to cover a large portion of the reciprocal space per crystal lamella at that resolution (Figure 3, video). For the fastest MicroED movie, the accumulated exposure was less than 2 e^-^ Å^-2^ total dose.

**Figure 3.**
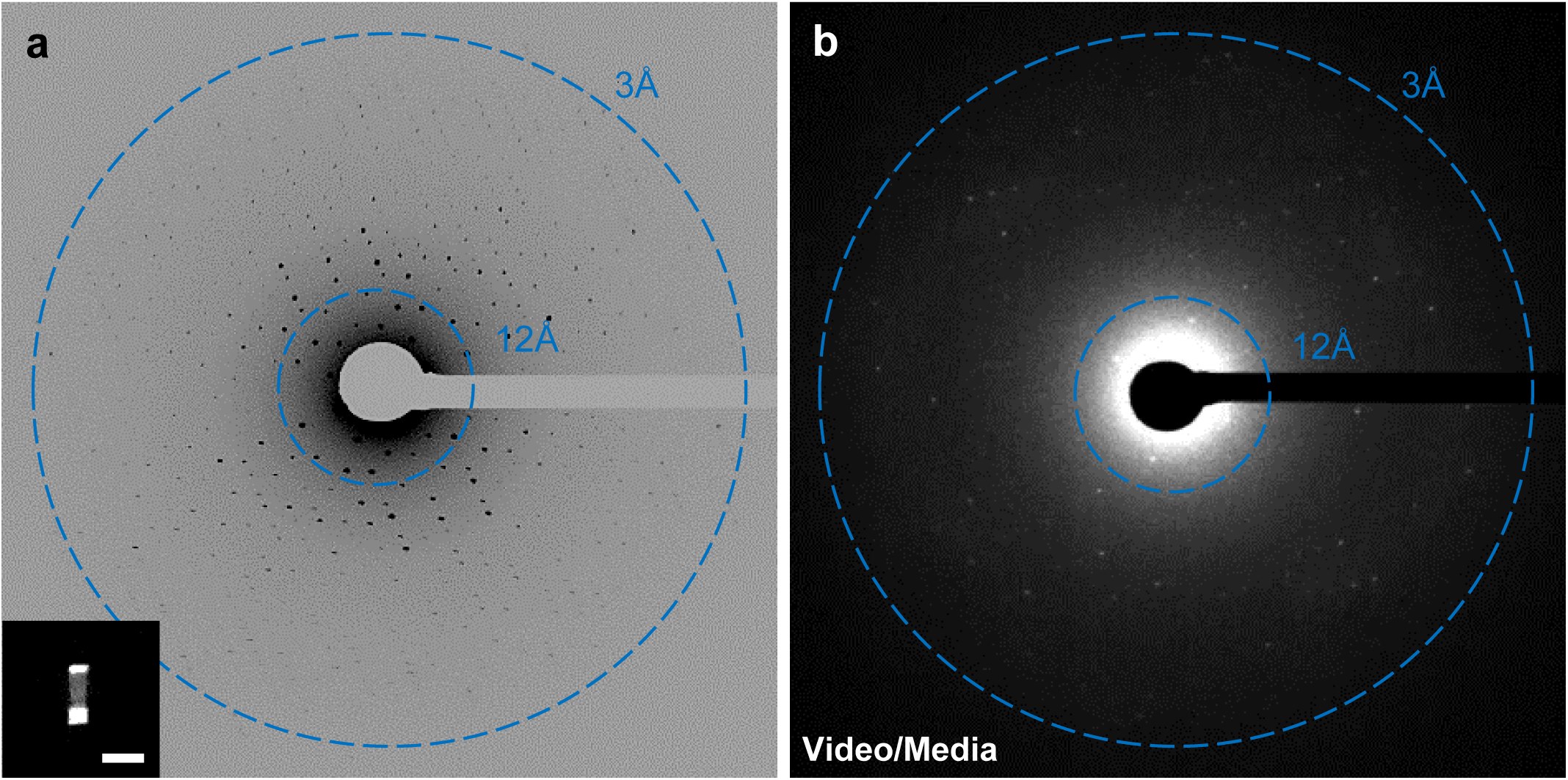
MicroED data collection from a milled mVDAC lamella. (a) A maximum intensity projection through 10 degrees of a MicroED dataset. Strong reflections are easily observed to ∼3A resolution. Inset: TEM micrograph of crystal lamella used for the data shown. Scale bar is 10um. (b) Movie of a mVDAC MicroED dataset from the same crystal as in (a).

The continuous rotation MicroED data were saved as MRC files and converted to SMV format using an in-house program that is freely available (https://cryoem.ucla.edu/MicroED). Images were processed in XDS (Kabsch, 2010a), with the negative pixels being lifted using a pedestal of 512 as previously described (Hattne et al., 2016, 2015). The space group was identified to be C 1 2 1 (#5), with a unit cell of (a, b, c) = (98.9 58.21 69.54), and (α, β, γ) = (90, 99.44, 90), which was very similar to the crystallographic parameters of the wild type mVDAC (Ujwal et al., 2008). A resolution cutoff was applied at 3.1Å, with an overall completeness of 80% (Table 1). Merging data from additional crystals did not increase the completeness because the mVDAC crystals have a preferential orientation on the grid. This observation was made previously using catalase crystals which also formed flat plates and had a preferential orientation (Nannenga et al., 2014a). The structure of mVDAC was solved by molecular replacement using the wild type model of VDAC (PDB accession code 3EMN). A single solution was found with a TFZ and LLG of 19 and 660, respectively. Following molecular replacement, the structure was inspected in COOT (Emsley and Cowtan, 2004) and refined.

Refinement of mVDAC followed standard procedures (Hattne et al., 2015; Shi et al., 2016). The model from molecular replacement was refined using electron scattering factors in PHENIX (Afonine et al., 2012). Model building and refinement were done iteratively until the refinement converged. The resulting R_work_ and R_free_ were 25% and 28%, respectively. The final model of mVDAC contains an N-terminal α-helix surrounded by a bundle of 19 β-sheets forming a barrel that encloses a hydrophilic pore (Figure 4a, 4b, video). This structure is similar to the wild type mVDAC with an all atom RMSD of 0.65 Å^2^.

**Figure 4.**
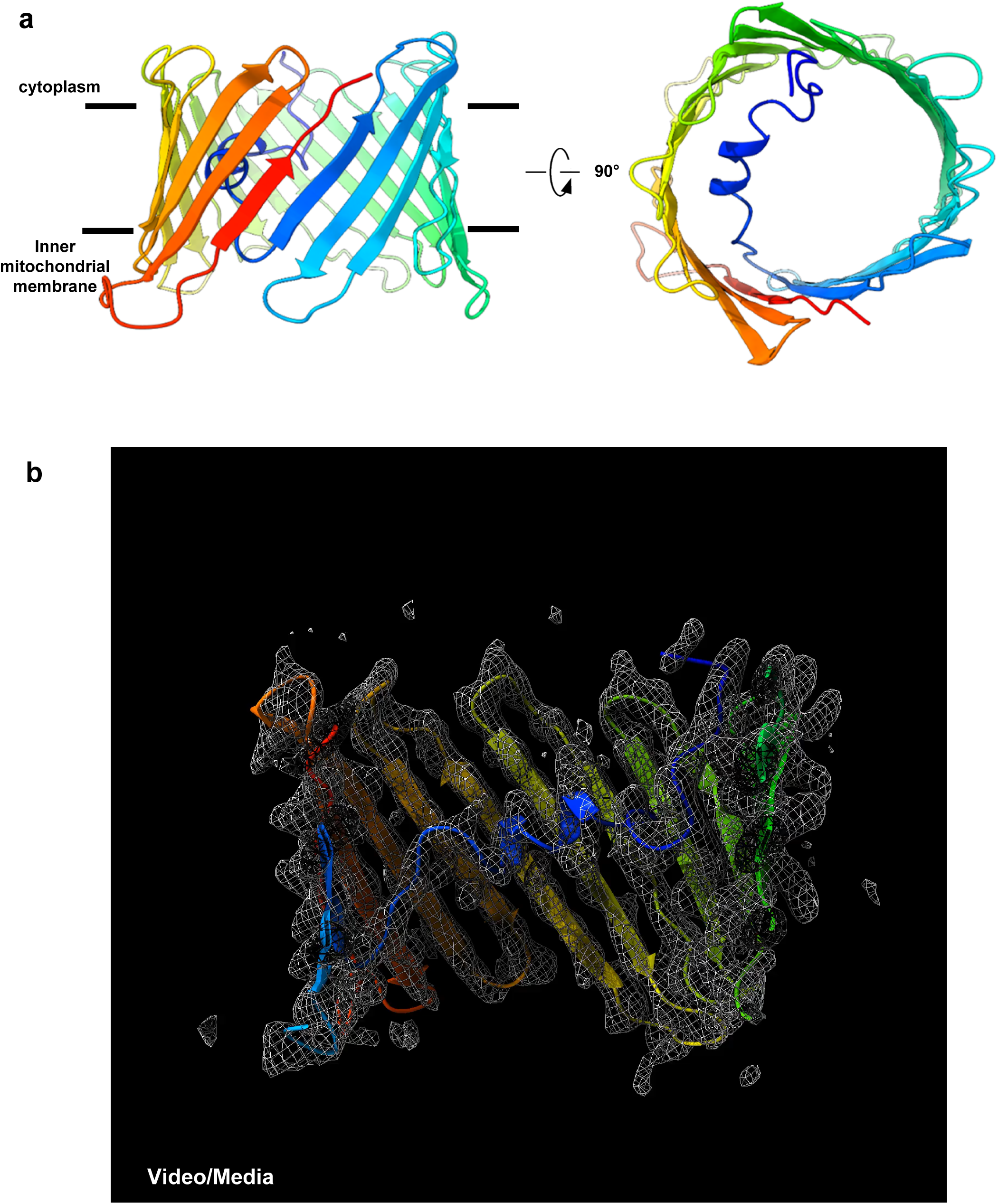
The structure of mVDAC by MicroED. (a) The final structure of mVDAC at 3.1A resolution in side and top views (left and right respectively). (b) Video showing the 2F_o_-F_c_ map of the entire protein through an entire rotation.

The packing of mVDAC monomers within the crystal lattice involved direct protein-protein contacts as well as contacts mediated by lipids. The different contact sites exist because individual mVDAC barrels do not pack as a planar hexagonal lattice as one would predict for a round monomer. Instead, each monomer makes a close contact with three monomers, and more distant contacts with an additional four monomers in a planar arrangement (Figure 5a). The three nearest neighbors of an mVDAC monomer are close enough for direct protein-protein crystal contacts but the other 4 neighboring barrels are too far, separated by up to 34 Å from one another. The spaces in between these distant mVDAC barrels are filled with lipids which act as glue to mediate crystal contacts (Figure 5b). We chose not to model the lipids at this stage because at this resolution we only observed discontinuous density for the lipid moieties. Lipids have been observed to mediate crystal contacts in other membrane protein studies. For example, the structure of the aquaporin-0 mediated membrane junction that was determined by electron diffraction from double layered crystals (Gonen et al., 2005, 2004). Lipids that mediated the crystal contacts were only observed once the resolution reached atomic resolution (Gonen et al., 2005) but were otherwise left unmodelled at the initial 3Å resolution study as the lipid densities were likewise fragmented (Gonen et al., 2004).

**Figure 5.**
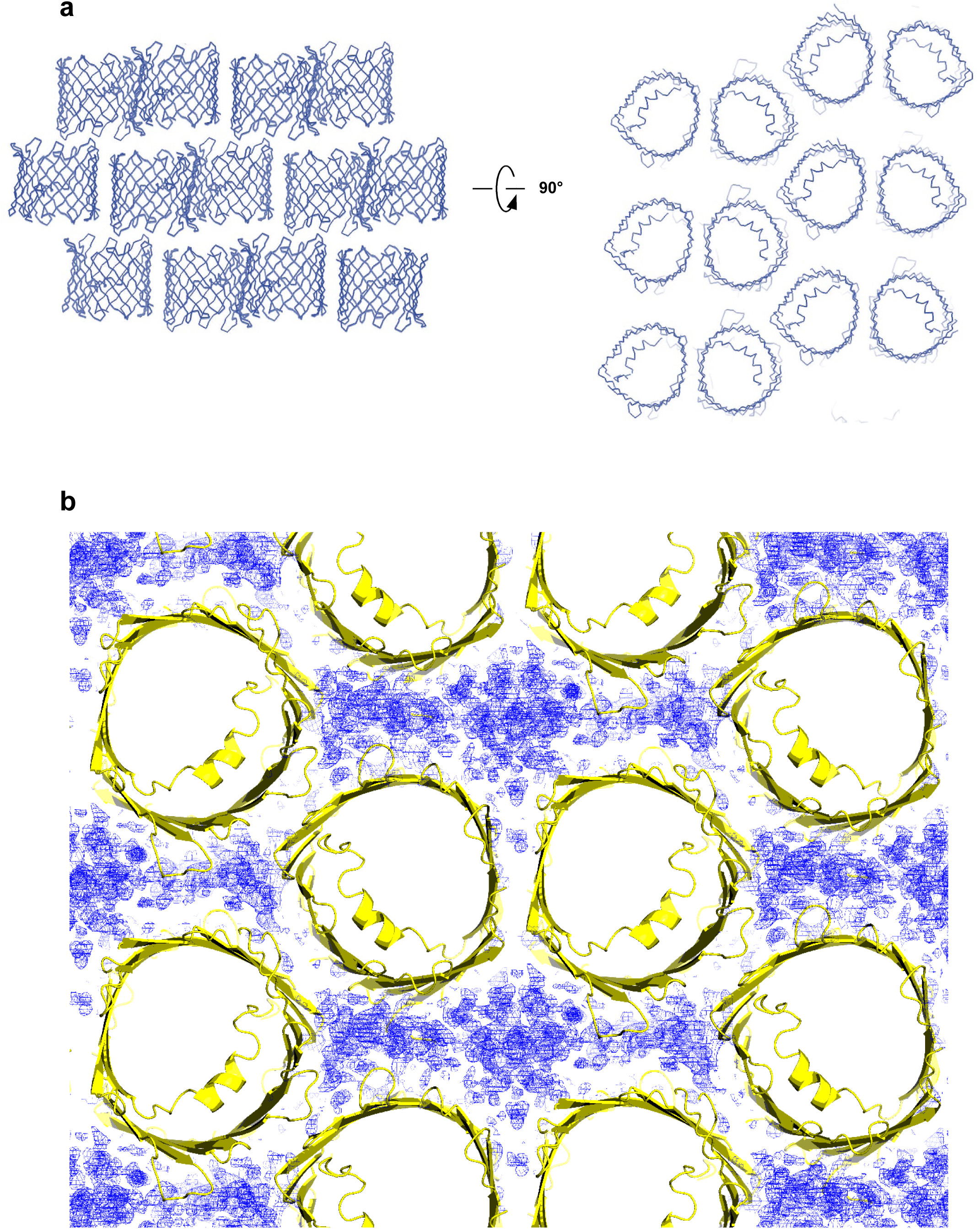
Crystal packing and contacts in mVDAC. (a) the lattice arrangement of the mVDAC molecules in the crystal in top and side views (left and right, respectively). (b) The 2F_o_-F_c_ map in blue between of the protein (yellow) is consistent with lipid molecules that likely mediate crystal contacts.

## Concluding Remarks

We have demonstrated a structure solution of a mammalian membrane protein grown in a modified lipidic environment that was not tractable by other crystallographic methods. While wild type mVDAC readily grows large crystals suitable for X-ray crystallography, the crystals here were barely visible and not amenable to analysis by X-ray crystallography. These crystals grew in a thick modified lipid matrix that made isolating crystals for MicroED challenging. In this case, FIB milling was necessary to remove excess material and allow structure determination by MicroED once crystal hydration was maintained during grid preparation. The MicroED structure was solved from a single crystalline volume of less than 1µm^3^, which is not possible using synchrotron X-ray crystallography. Finally, as future studies will focus on improving the resolution of this approach we expect to fully resolve the lipids that mediate crystal contacts in these, and, in the future, other crystals of membrane proteins grown in a lipid matrix and studied by MicroED. This combination of methodologies could open the door to investigating the influence of lipids on membrane protein structure and function in ways that were previously not possible.

## Materials and Methods

### Contact for reagent and resource sharing

Further information and requests for resources and reagents should be directed to and will be fulfilled by the lead contact, Tamir Gonen.

### Protein production and purification

Lipid bicelles were prepared as described before (Faham and Bowie, 2002). Mutant VDAC was expressed and purified as described (Ujwal et al., 2008). Purified mVDAC was concentrated to 15 mg/ml, and mixed in a 4:1 protein/bicelle ratio with a modified bicellar solution, resulting in 12 mg/ml mVDAC1 in 7% bicelles.

### Protein crystallization

Crystal screens for mVDAC started from the known crystallization conditions conducted at 20°C (Ujwal et al., 2008) and modified for crystal growth. Microcrystals appeared in a well containing 20% MPD, 0.1 M Tris·HCl (pH 8.5) with 10% PEG400 added to the protein drop only.

### Grid preparation

Quantifoil R2/2 Cu200 grids were glow discharged for 30s prior to use. The blotting chamber was set to 18°C and 90% humidity, and filter paper was added. The system was allowed to equilibrate for 15 mins before any blotting was conducted. 2µL of mother liquid was added to the grid inside of a vitrification robot to prevent any dehydration during application. A small portion (∼0.5µL) from the central lipid clump within the crystallization drop was pipetted carefully into the already applied 2µL of mother liquor. The sample chamber of the vitrification robot was held at 90% humidity. This mixture was allowed to incubate for 20s, and was then gently blotted from the back and immediately vitrified in liquid ethane as described (Martynowycz and Gonen, 2020). Grids were transferred and stored in liquid nitrogen prior to further investigation.

### Cryo-SEM imaging and Cryo-FIB milling of VDAC

All FIB/SEM experiments were performed on a Thermo-Fisher aquilos dual-beam FIB/SEM instrument at liquid nitrogen temperatures as described (Duyvesteyn et al., 2018; Martynowycz et al., 2019a, 2019b; Wolff et al., 2020). The instrument was operated at 2kV and 3.1pA while using the SEM for imaging, and 30kV and 1.5 or 10pA while operating the FIB beam for imaging. The grid was sputter coated in platinum for ∼1min using a high current in order to apply a 500nm thick layer of platinum evenly over the entire grid. An all-grid map was collected in the MAPS software (Thermo-Fisher), where potential crystals were identified by looking for sharp edges in areas not over the grid bars. Each grid square was individually inspected again using the FIB at 1.5-10pA current to verify the already selected crystals or identify new crystals. Verified mVDAC crystals were brought to eucentric height prior to milling. Milling was conducted as described. In general, milling was done in batch, where each step of rough, fine, and ultra-fine (or, polishing) were conducted on each crystal prior to additional thinning. In this way, contamination of the lamellae by amorphous ice was minimized.

### MicroED data collection

Data collection was performed as described(Hattne et al., 2015; Martynowycz et al., 2019a; Nannenga et al., 2014b; Shi et al., 2013). MicroED was tested on a 200kV Talos Arctica equipped with a Ceta-D CMOS. Individual diffraction patterns were collected from each lamella without rotation to evaluate the quality of diffraction that could be obtained. Data were collected on a cryogenically cooled Titan Krios operating at 300kV, corresponding to an electron wavelength of 0.0197Å. Lamellae were identified in low-magnification montage taken at approximately 64x magnification on a Ceta-D camera (Thermo-Fisher). Data were collected under continuous rotation from the three lamellae at rotation speeds between 0.1-0.3 ° s^-1^ with frames being read out at every 5, 3, and 2s. The general strategy was to collect data with individual wedges corresponding to approximately 0.5°. The exposure rate was set to less than 0.01 e^-^ Å^-2^ s^-1^ to limit radiation damage (Hattne et al., 2018). Data were recorded using a selected area aperture that limited the exposed area to region approximately 2µm in diameter. Since the lamella thickness was merely ∼200nm the MicroED data was collected from a crystalline volume of less than 1μm^3^. Data were saved as single MRC stacks prior to data analysis.

### MicroED data processing

Data were converted from MRC to SMV format using software that is freely available (https://cryoem.ucla.edu/). Converted frames were used to index, integrate, and scale the collected in XDS and XSCALE (Kabsch, 2010a, 2010b). Individual datasets were optimized by removing frames from the end to improve the scaling between the three datasets. A resolution cutoff was applied at 3.1Å, where the CC_1/2_ of the merged data remained positive. Further reduction of the resolution could be done during refinement based upon the refinement statistics. Molecular replacement was performed in PHASER (McCoy et al., 2007) using the PDB search model 3EMN (Ujwal et al., 2008) and the merged intensities. A single solution was identified using PHASER in space group C 1 2 1, or #5. The initial model from PHASER was inspected in coot (Emsley and Cowtan, 2004) prior to refinement. Refinement was conducted by PHENIX.REFINE using electron scattering factors (Adams et al., 2011). We found that the best statistics were obtained by not reducing the resolution further, since the outermost shell of our refinement (3.21 −3.12) did not have a particularly poor R_free_ value (0.36). The structure was left as-is without further modelling any densities outside of the protein residues, as these are to be detailed in future work regarding the functional importance of this mutation.

### Figure preparation

Figures were arranged in Microsoft Powerpoint and Photoshop 2020; Chimera X and coot. Images were adjusted and cropped in FIJI (Schindelin et al., 2012). Tables were arranged in Microsoft Excel. Protein models and meshes were generated using PyMol (*The PyMOL molecular graphics system*, 2014). Movies of the density map and structure were recorded in Chimera X.

## Acknowledgements

This study was supported by the National Institutes of Health P41GM136508 to T.G. and R35 GM135175 J.A.. The Gonen lab is supported by funds from the Howard Hughes Medical Institute. The structure factors and coordinates are deposited in the PDB and the associated map in the EMDB.

## Author contributions

FK purified and crystallized mVDAC in modified bicelles. MWM prepared cryoTEM grids, did FIB milling and collected MicroED data. The structure of mVDAC was solved by MWM and FK. Figures were prepared by JH, TG and MWM. All authors participated in writing and approving the manuscript. The project was conceived by JA and TG.

## Figure legends and tables

**Table 1. MicroED Crystallographic Table mVDAC.**

---

**Figure S1:**
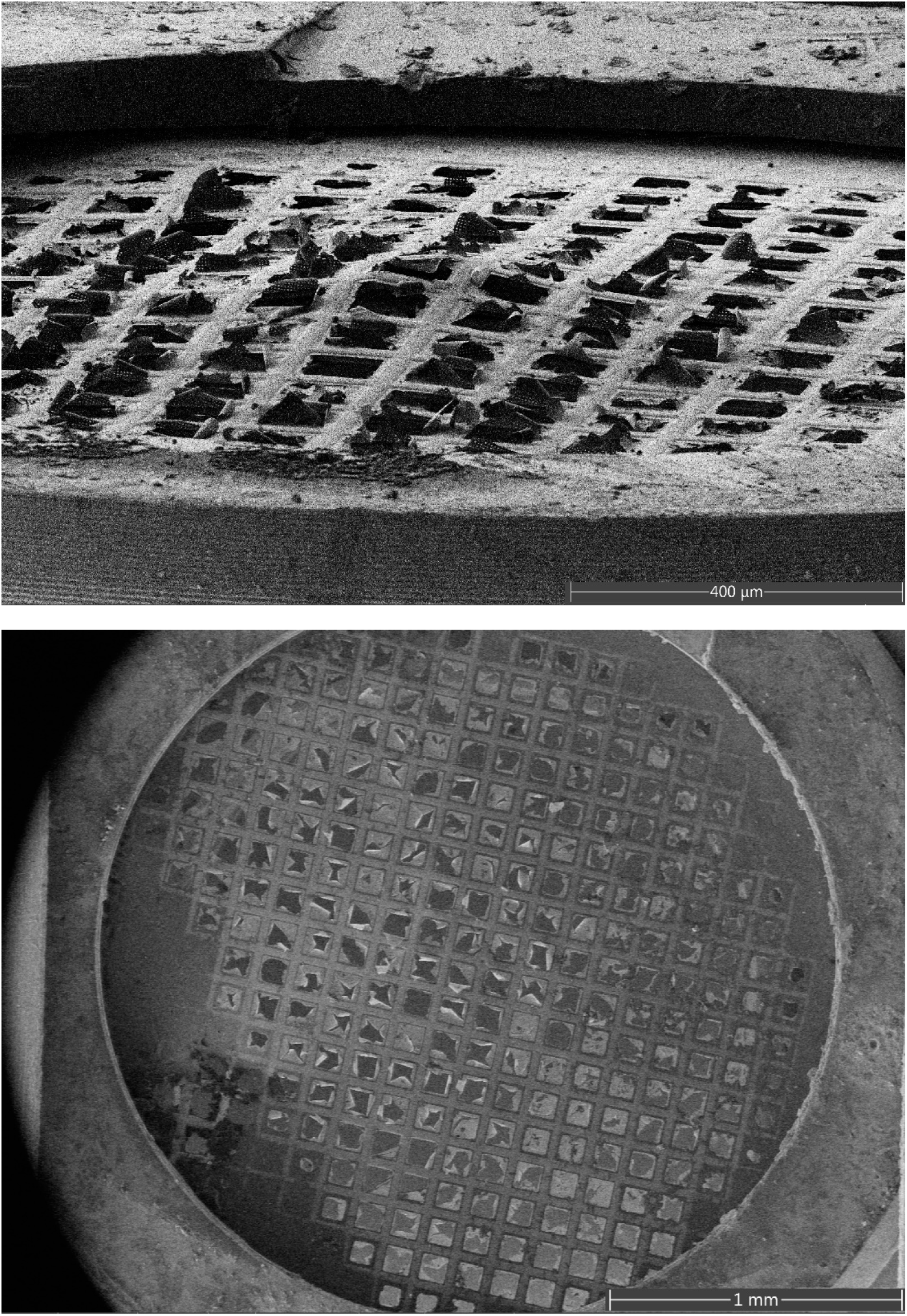
The grid was blotted in a vitrobot from both the front and back. Most windows were broken, and no usable areas were identified

**Figure S2:**
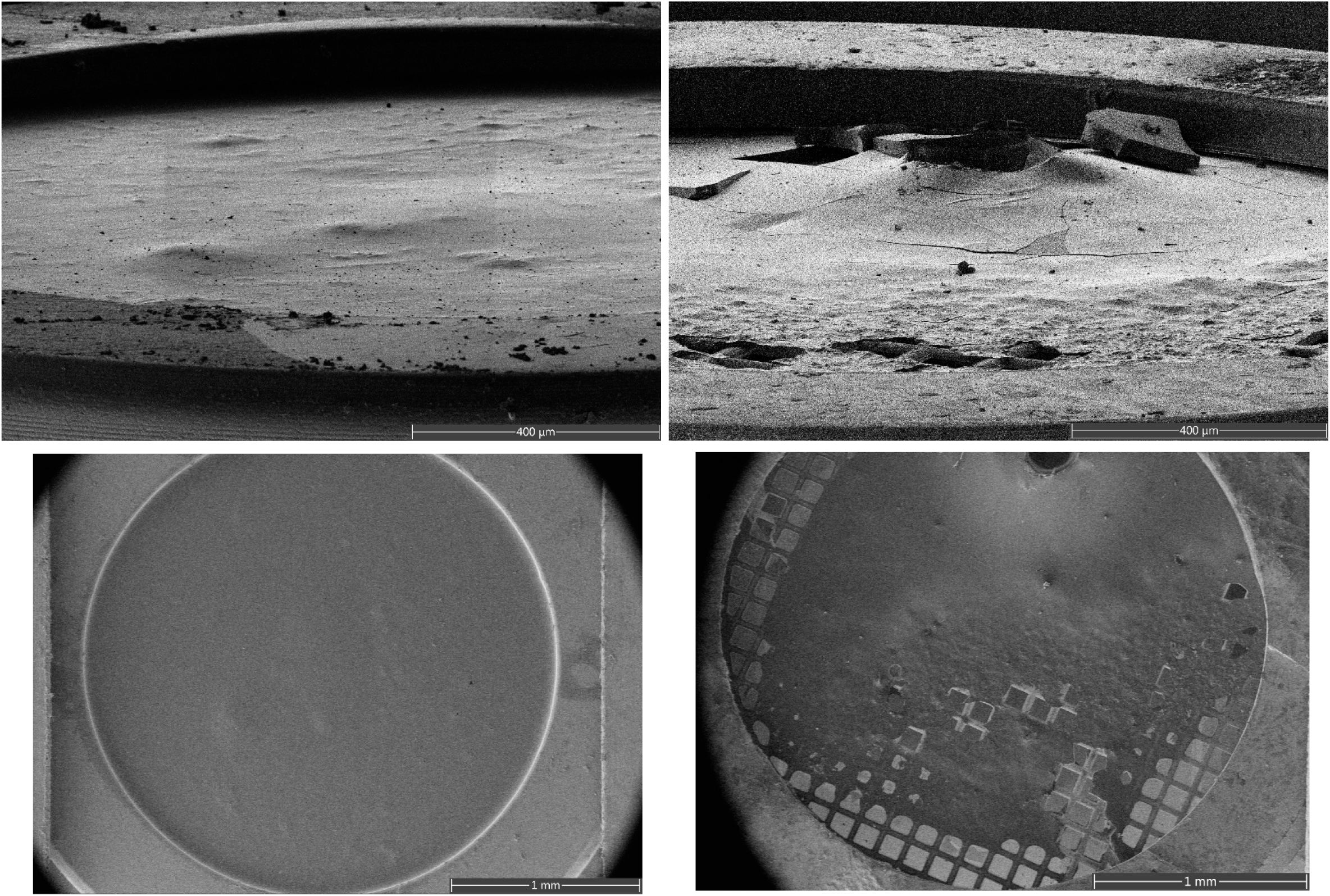
The entire viscous crystallization drop was pipettes onto the grid. No rinsing and blotted from the back only. A uniform thick layer of material was observed by no areas that clearly appeared to contain crystals identified.

**Figure S3:**
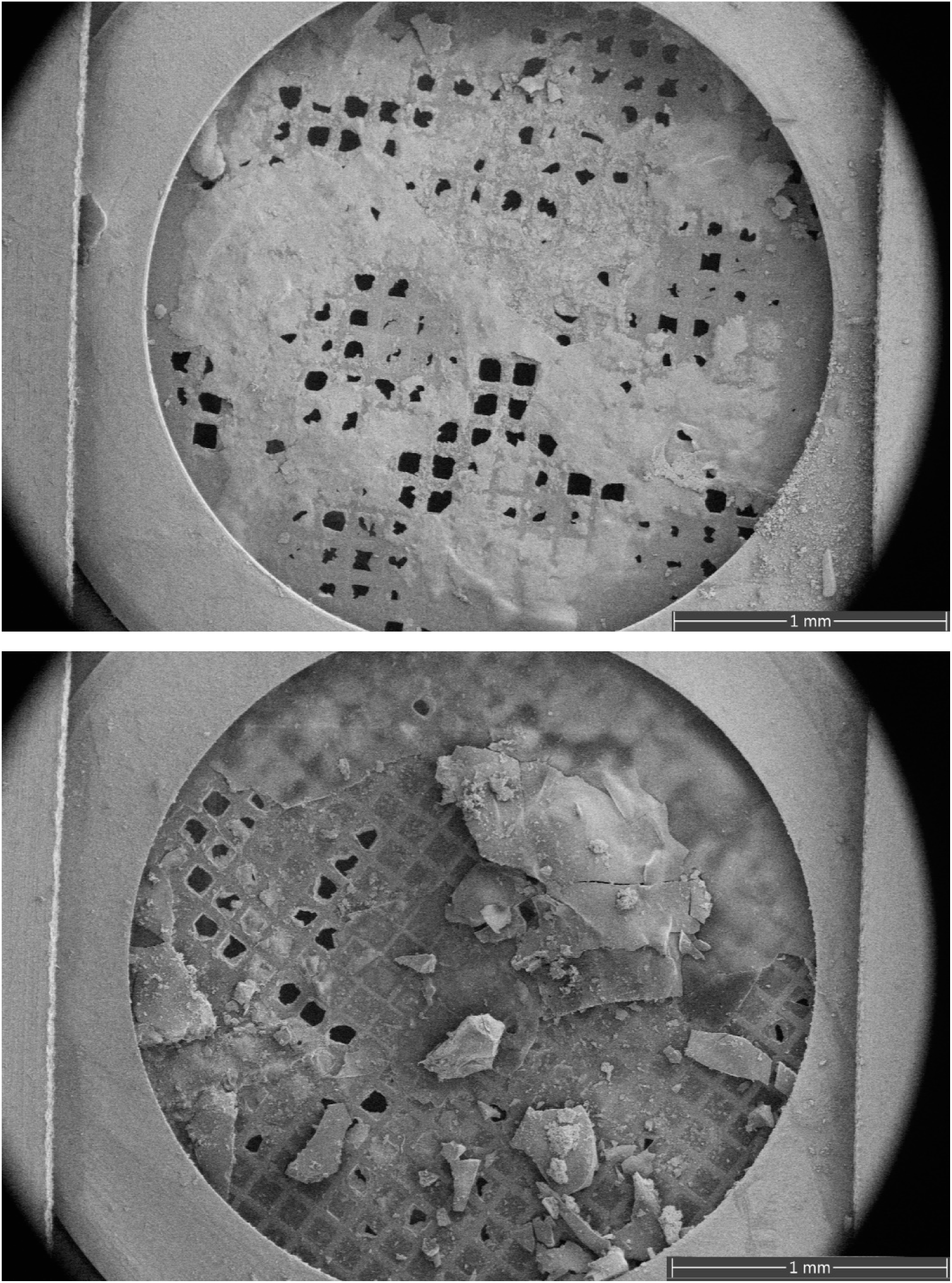
The entire viscous crystallization drop pipettes onto the grid and blotted from the front only and plunged into liquid nitrogen. Many windows appeared broken while the majority of the grid was covered in thick areas of ice.

**Figure S4:**
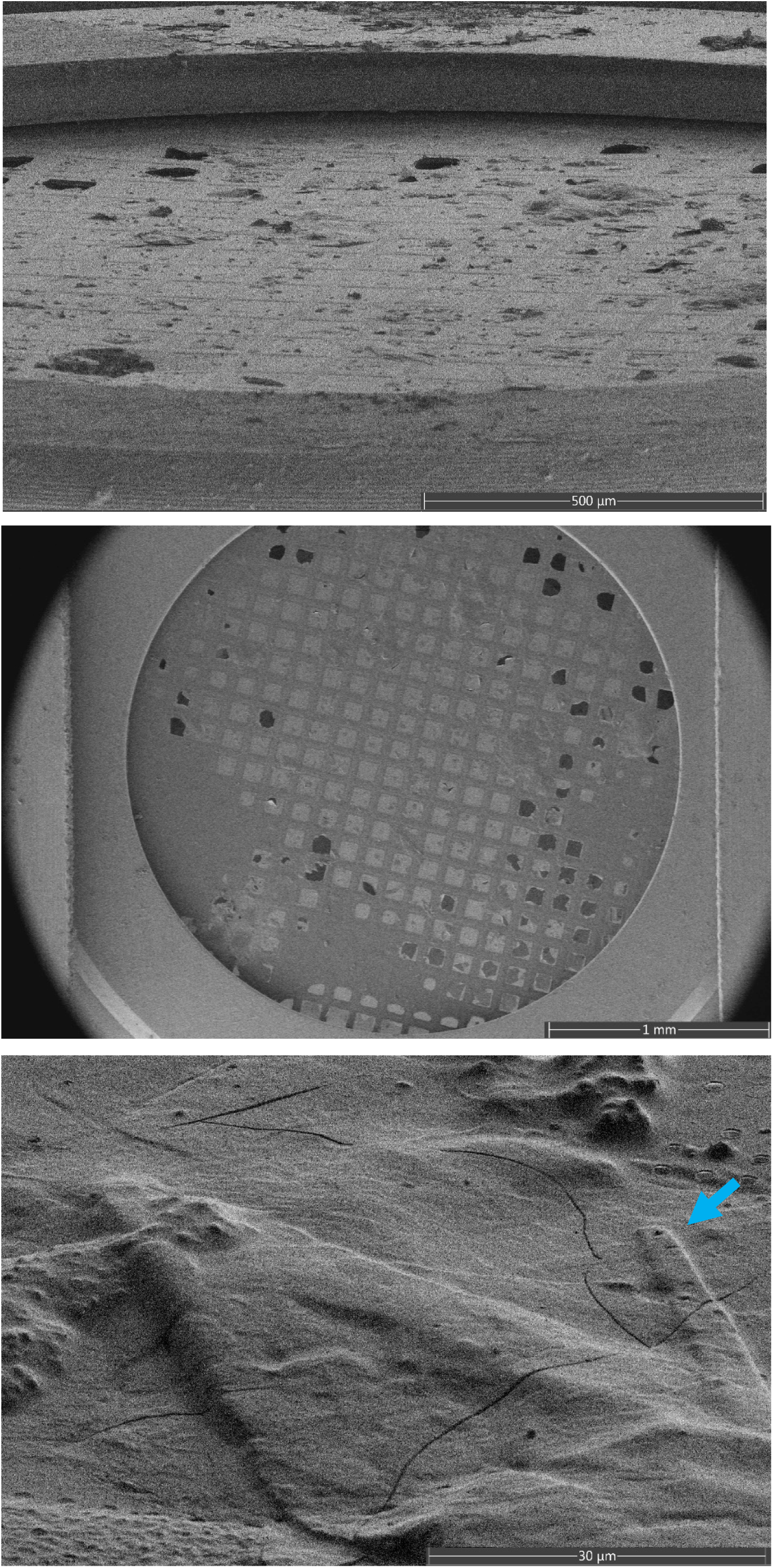
The viscous crystallization drop was applied to the grid with low humidity and blotted from the back before plunging into ethane. Most windows appeared intact and some areas that may be crystalline were identified (arrow). The combination of back blotting and ethane improved the preparation. No data could be obtained from this grid.

**Figure S5:**
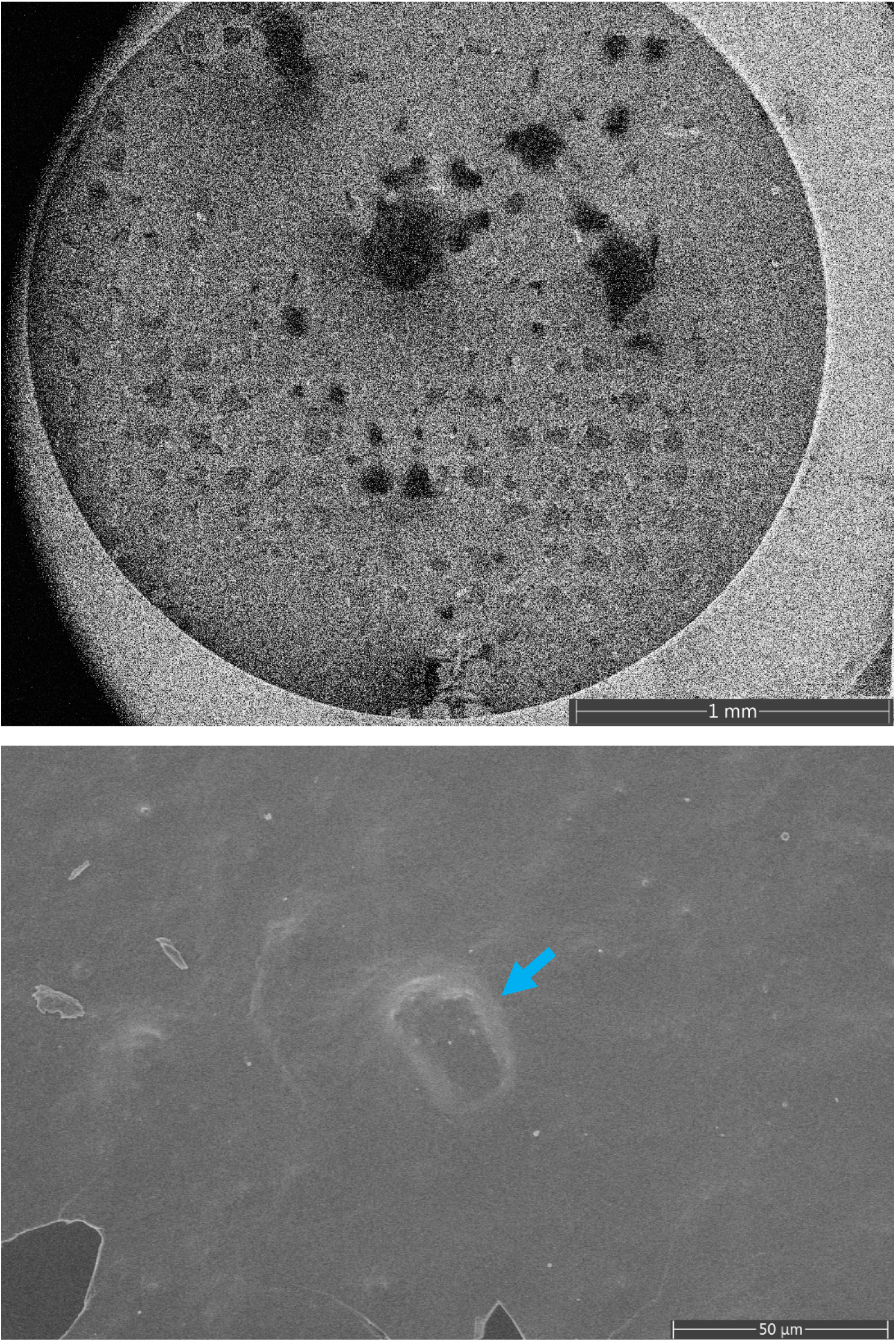
The viscous crystallization drop was applied to the pre wetted grid. The vitrobot was set to low humidity and at room temperature and blotting was done from the back followed by plunging into ethane. The grid was covered uniformely with the crystalline solution and at high magnification items consistent with crystals could be identified (arrow). This preparation yielded low resolution data (∼8A)

**Figure S6:**
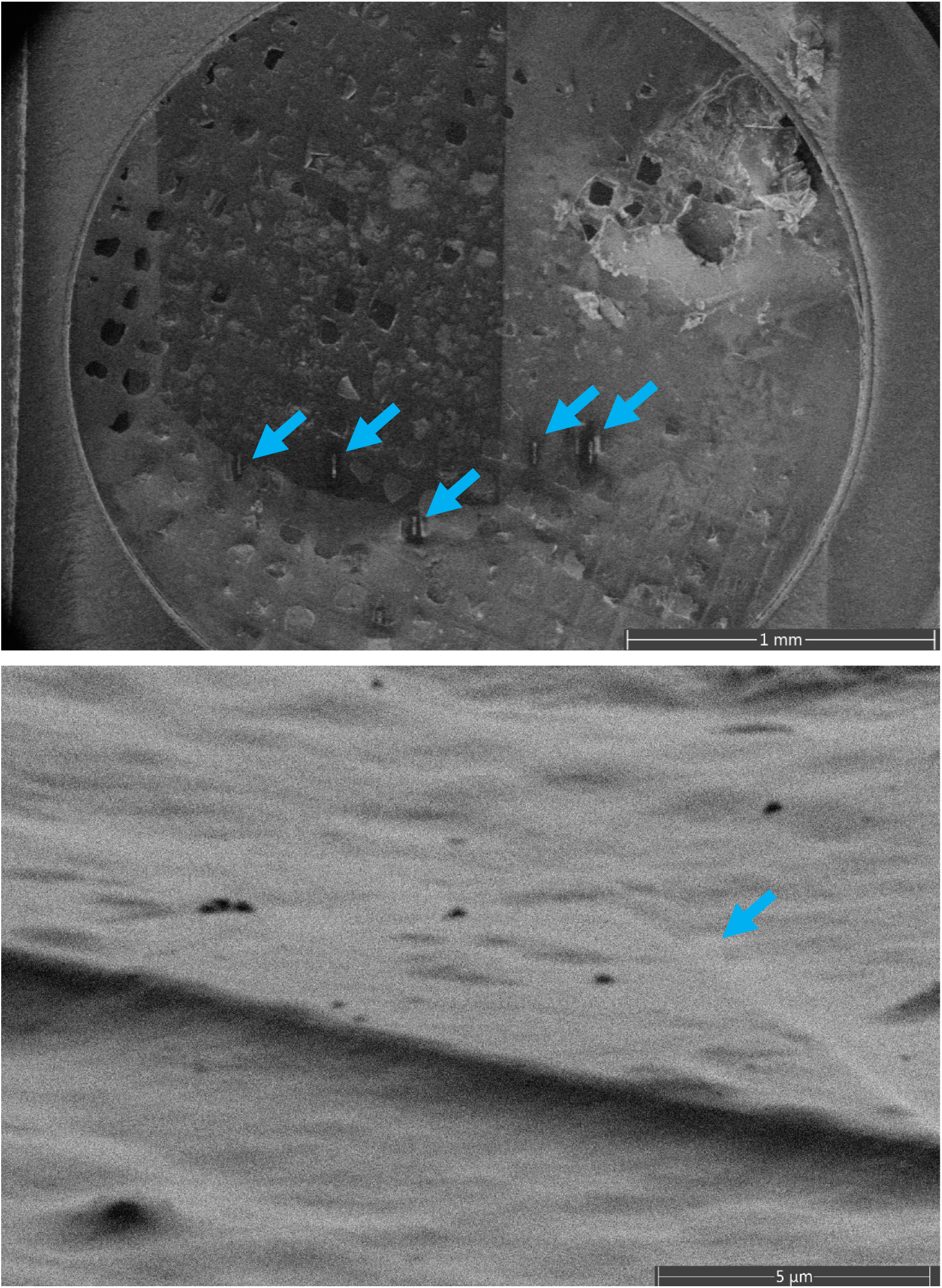
The viscous crystallization drop was applied to the pre wetted grid. The vitrobot was set to low humidity and to 4 degrees C and blotting was done from the back followed by plunging into ethane. The grid was covered with the crystalline solution and at high magnification items consistent with crystals could be identified (arrow). This preparation yielded higher resolution data (∼4A)

**Figure S7:**
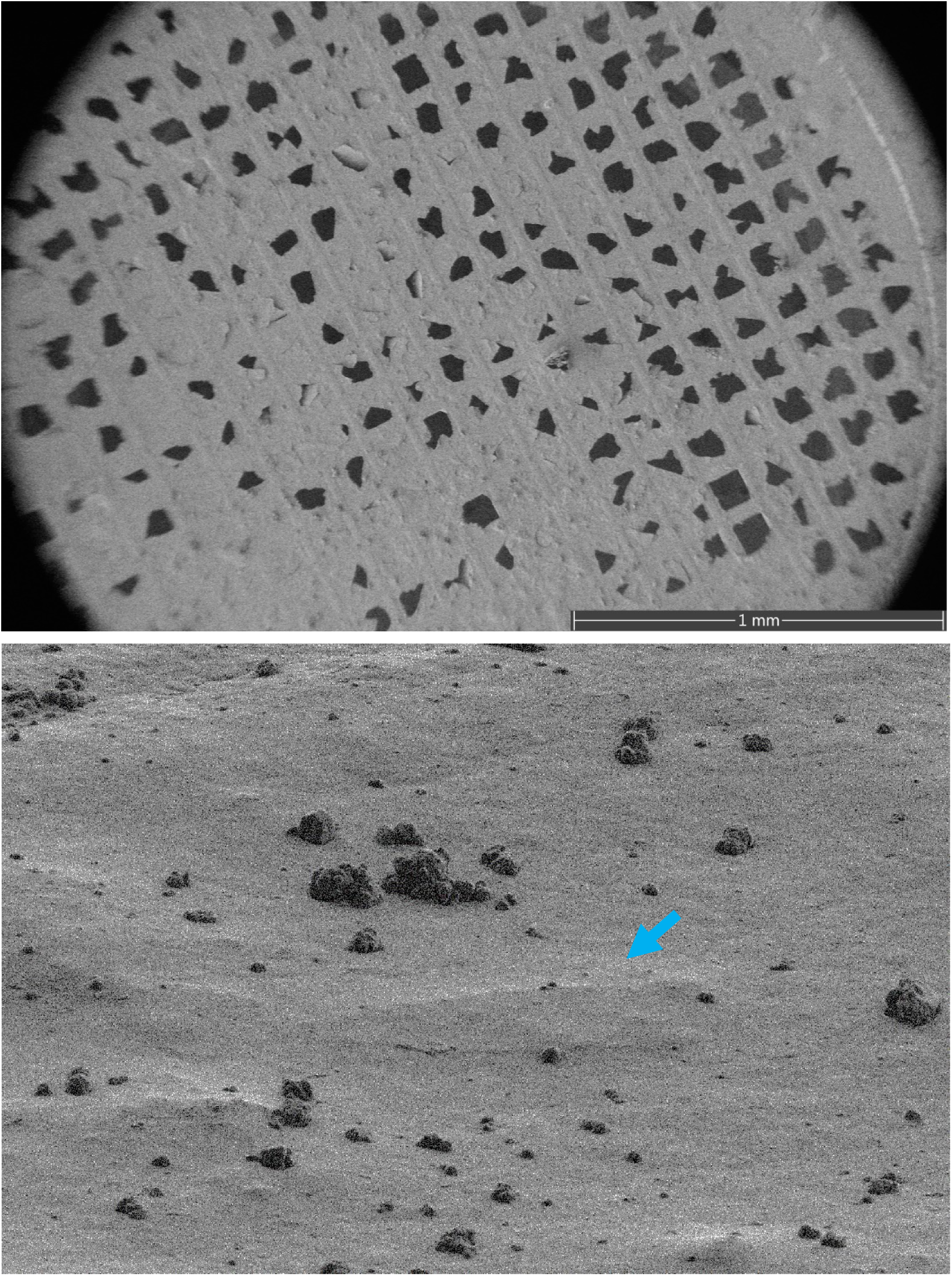
The viscous crystallization drop was applied to the pre wetted grid. The vitrobot was set to 90% humidity and to 4 degrees C and blotting was done from the back followed by plunging into ethane. Although some windows were broken, the grid had many intact windows that were covered with the crystalline solution and at high magnification items crystals could be identified (arrow). This preparation yielded the best data after milling (∼3A)

**Figure S8:**
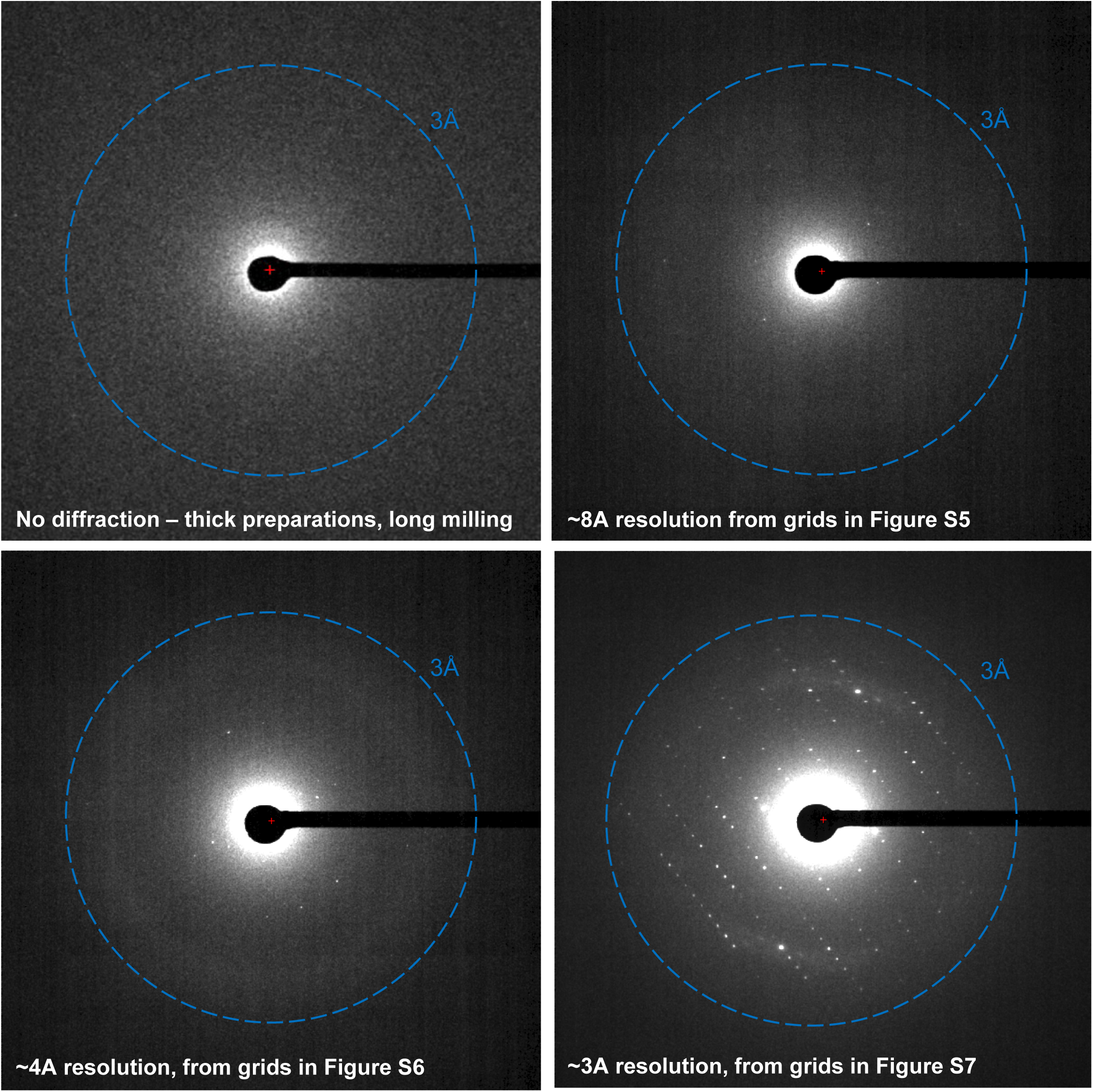
Progression in diffraction quality as grid preparation method improved. The best data at ∼3A resolution was obtained from grids that were prepared using pre-wet grids held at 90% humidity, 4 degrees C, back blotted and plunged into ethane.

**Figure S9:**
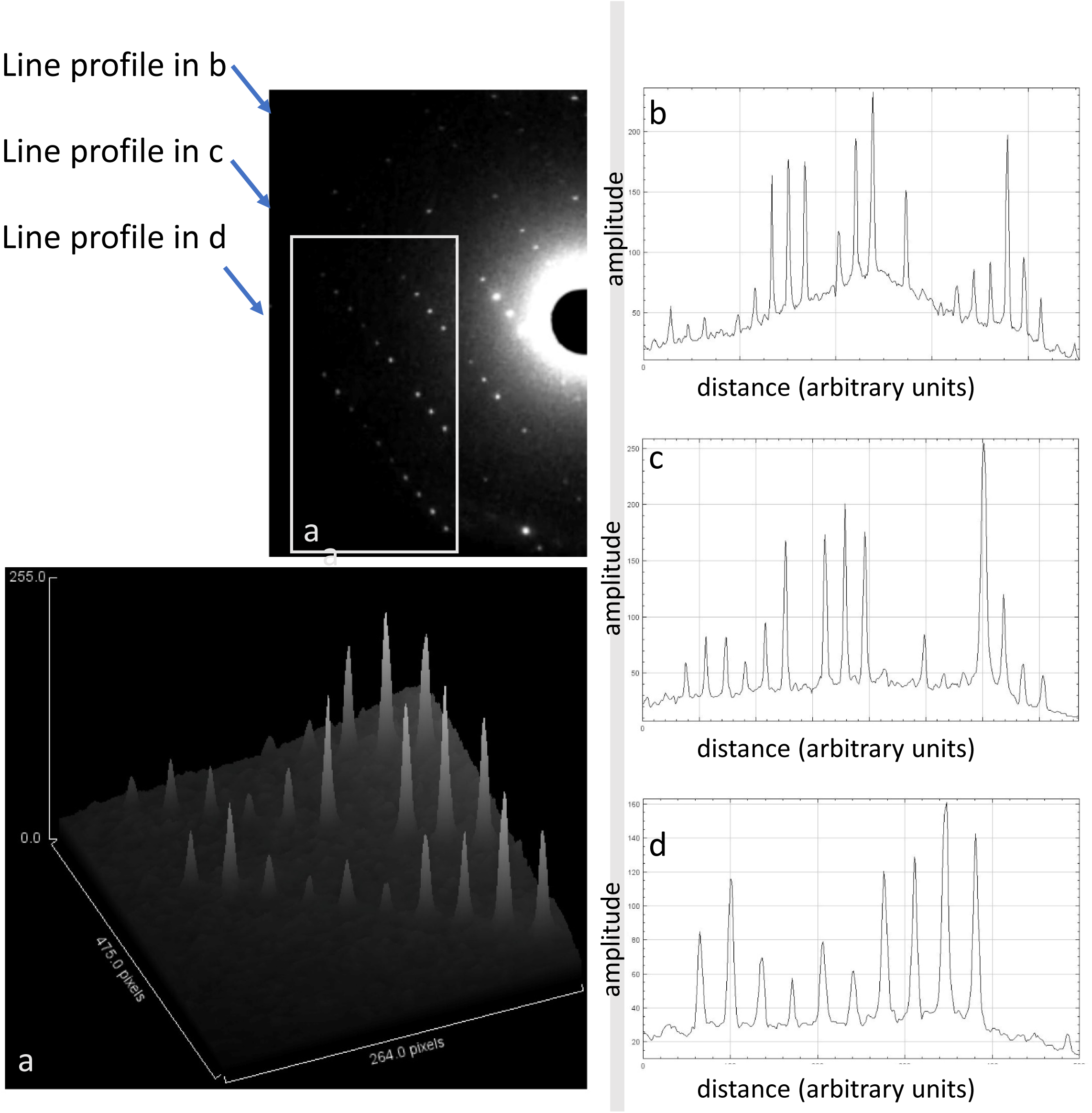
Quality of the MicroED data obtained from ∼200nm lamella of membrane protein crystals embedded in a thick lipid matrix. This data was obtained from the crystal shown in Figure 2 of the main text

